# Neutralisation of SARS-CoV-2 Omicron subvariants BA.2.86 and EG.5.1 by antibodies induced by earlier infection or vaccination

**DOI:** 10.1101/2023.10.01.560365

**Authors:** Ria Lassaunière, Charlotta Polacek, Sharmin Baig, Kirsten Ellegaard, Leandro André Escobar-Herrera, Anders Fomsgaard, Katja Spiess, Olivier Schwartz, Delphine Planas, Etienne Simon-Lorière, Uffe Vest Schneider, Raphael Niklaus Sieber, Marc Stegger, Tyra Grove Krause, Henrik Ullum, Pikka Jokelainen, Morten Rasmussen

## Abstract

Highly mutated SARS-CoV-2 Omicron subvariant BA.2.86 emerged in July 2023. We investigated the neutralisation of isolated virus by antibodies induced by earlier infection or vaccination. The neutralisation titres for BA.2.86 were comparable to those for XBB.1 and EG.5.1, by antibodies induced by XBB.1.5 or BA.4/5 breakthrough infection or BA.4/5 vaccination.

## Introduction

In late July 2023, global SARS-CoV-2 surveillance programs identified a new, highly mutated Omicron subvariant, BA.2.86. This variant is a descendant of the Omicron BA.2 variant, with the most recent common ancestor estimated to be from between 9th April and 24th July 2023 (1,2). Since the emergence of BA.2.86, multiple countries have reported its presence, primarily as sporadic BA.2.86 cases with no clear epidemiological link. From the first 12 BA.2.86 cases in Denmark, the estimated effective reproduction number (R_e_) of this variant was 1.29-fold greater than that of XBB.1.5 and at least comparable to that of EG.5.1, one of the most rapidly expanding XBB subvariants (3). This is consistent with an outbreak of BA.2.86 in a care home in the United Kingdom (UK) that experienced an attack rate above 85%, indicating high transmissibility in close contact situations (4).

The earliest cases of BA.2.86 in Denmark suggested a clinical presentation and disease severity no different from other SARS-CoV-2 variants circulating during July/August 2023, such as EG.5.1 and XBB.1.16 (1). Symptoms included cough, shortness of breath, and fever; none were severely ill. Some of the early cases had an underlying disease or received immune-modulating treatment. Taken together with the care home outbreak in the UK (4), individuals of older age and with comorbidities appear to be more at risk of developing a symptomatic BA.2.86 infection that warrants medical attention.

The increasing numbers of BA.2.86 infections 1-2 years after 3-4 immunisations, with or without prior infection, suggests that waning immunity may contribute to the susceptibility to BA.2.86 infection. Moreover, the highly mutated spike protein of BA.2.86 is antigenically distinct from all other SARS-CoV-2 variants, including the XBB subvariants, as determined using mRNA-vaccinated mouse serum (5). This suggests that BA.2.86 may evade pre-existing humoral immunity, as well as the acquired immunity from the updated XBB.1.5-based vaccines. The care home outbreak data suggested limited vaccine effectiveness against infection with BA.2.86 four months after vaccination (4).

To estimate the cross-neutralisation of BA.2.86 by XBB.1.5-induced antibodies, we used serum or plasma samples from persons who had experienced an XBB.1.5 infection as a proxy for the humoral immune response generated by the updated XBB.1.5 vaccines. In contrast to assessing antibodies raised against single SARS-Cov-2 variants in vaccinated or infected animals (5), human XBB.1.5 breakthrough infections are more representative of the complex and diverse pre-existing immunity of future XBB.1.5-based vaccine recipients (6). As a representative of the 2022 SARS-CoV-2 variant vaccine, we assessed antibody responses in persons who had either experienced an Omicron BA.4/5 breakthrough infection or received the Comirnaty Original/Omicron BA.4-5 bivalent mRNA vaccine as a fourth vaccination.

## Results

SARS-CoV-2 variants exhibit variable affinities for the angiotensin-converting enzyme 2 (ACE2) host receptor and dominance in using either the fusion or the endocytic entry pathway in immortalised cell lines (6,7). To account for this, we attempted viral isolation on different cell lines permissive to SARS-CoV-2 infection with respiratory sample material obtained from individuals with an RT-PCR- and sequencing-confirmed BA.2.86 infection. Using a protocol detailed in the Method section, BA.2.86 established a primary infection from clinical sample material in human-derived cell lines CaLu-3 and iGROV-1, but not African Green Monkey kidney-derived cell lines Vero E6, Vero/hSLAM, and VeroE6 expressing ACE-2 and TMPRSS2. In contrast, EG.5.1 was readily isolated from Vero E6 cells. BA.2.86 infection of CaLu-3 and iGROV-1 cells from a single clinical sample did not yield cell line-specific spike protein adaptations, as determined by sequencing of these cell culture progenies. Once isolated, BA.2.86 efficiently infected Vero E6 cells and induced a clear cytopathic effect (**Fig S1**).

The spike protein of BA.2.86 contains approximately 60 amino acid changes relative to the SARS-CoV-2 index strain (**Fig 1A**). To assess its potential immune evasiveness, we first determined neutralising antibody titres in a live virus neutralisation assay for 13 individuals who experienced an XBB.1.5 breakthrough infection of whom the majority (n=12) received at least three prior vaccinations. The geometric mean neutralising antibody titre to D614G (B.1), BA.2, BA.5, XBB.1, EG.5.1 and BA.2.86 were 376, 363, 464, 99, 123, and 61, respectively (**Fig 1B**). The neutralisation titres for XBB.1, EG.5.1 and BA.2.86 were 3- to 7.6-fold lower compared to the ancestral D614G strain and the Omicron BA.2 and BA.5 variants. The geometric mean titre for BA.2.86 did not differ significantly from that of XBB.1 and EG.5.1.

**Figure 1.**
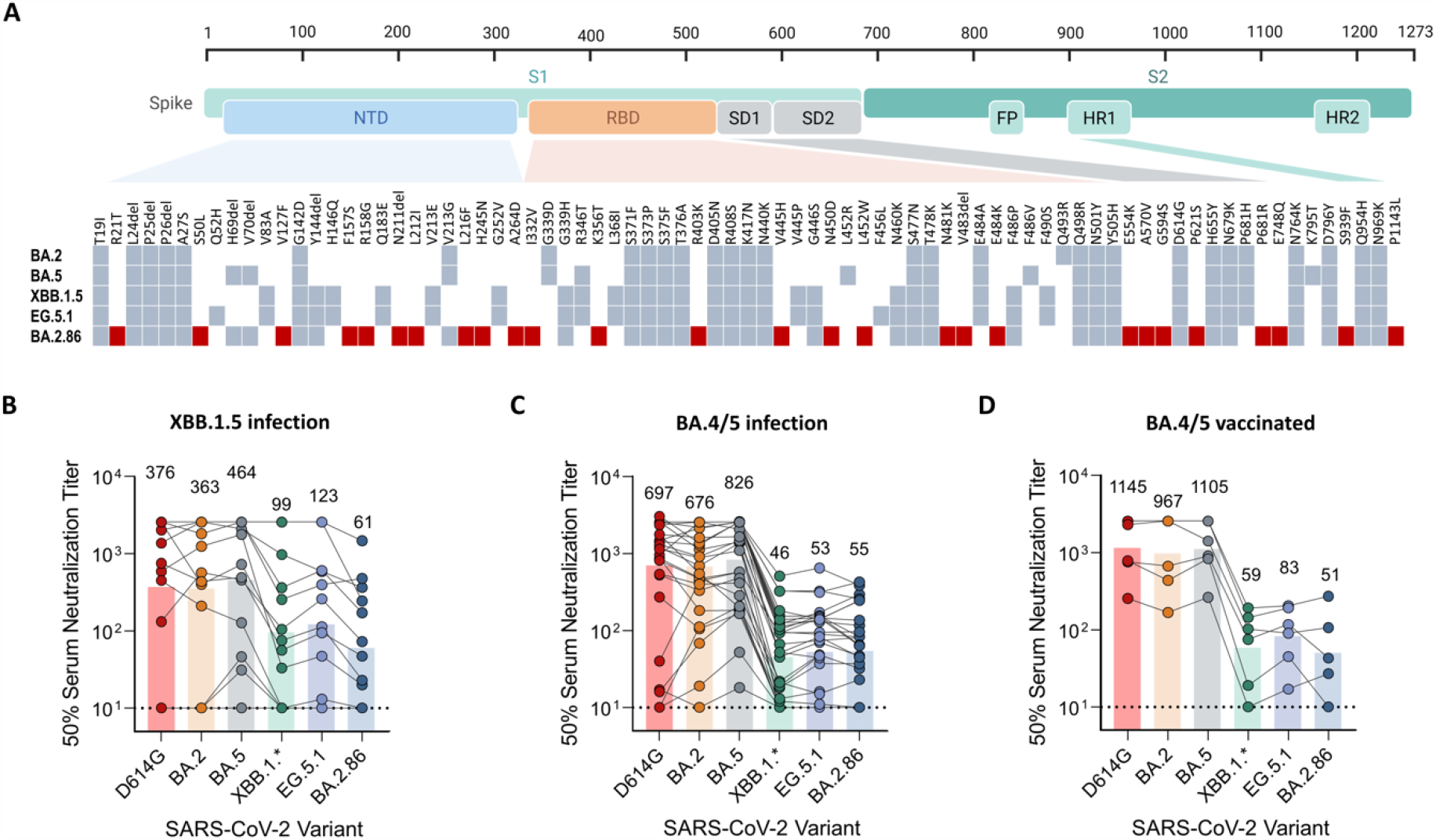
Immune evasion potential of BA.2.86 A) Spike mutations of SARS-CoV-2 variants BA.2, BA.5, XBB.1.5, EG.5.1, and BA.2.86 relative to the spike protein domains of SARS-CoV-2 index strain. Amino acid changes unique to BA.2.86 are indicated in red. B-D) Relative neutralising antibody levels for individuals following a breakthrough infection with an XBB.1.5 spike protein variant (B), a breakthrough infection with an Omicron BA.4/5 variant (C), and recipients of the bivalent Omicron BA.4/5 mRNA vaccine (D).

We next evaluated neutralising antibody titres for individuals who experienced a BA.4/5 breakthrough infection (n=30; of whom 23 received at least three prior index strain vaccinations) (**Fig 1C**). To compare we also assessed BA.2.86 virus neutralisation for individuals who had received the bivalent BA.4/5 mRNA vaccine as a fourth vaccination (n=6) (**Fig 1D**). For both these groups, the geometric mean titres to D614G, BA.2, and BA.5 were higher than those for individuals who experienced an XBB.1.5 infection and marginally lower to XBB.1, EG.5.1, and BA.2.86 compared to the same group. Similar to that observed for the XBB.1.5 infected group, the neutralisation titres for BA.2.86 did not differ significantly compared to titres for XBB.1 and EG.5.1.

## Discussion

The Omicron BA.2.86 subvariant has a distinctly different, highly mutated spike protein that alters its *in vitro* infectivity characteristics and antigenicity relative to earlier variants (5,6). This raises concern over the effectiveness of 2023/2024 updated vaccines derived from the Omicron XBB.1.5 spike protein. Data from this study and others suggest that, despite the highly different spike mutation constellation of BA.2.86 that alters monoclonal antibody specificities, the overall neutralisation ability of polyclonal serum antibodies after an XBB.1.5 breakthrough infection is comparable to that of circulating XBB recombinant strains (2,5,6,8–10).

Multiple factors relating to the virus, vaccine, and host modulate the antibody response in a person. These include vaccine modality, number of vaccine doses, time since vaccination, homologous or heterologous vaccination, the interval between doses, virus antigenicity, hybrid immunity (combination of prior infection and vaccination), immune imprinting, host age, sex, and immune-modulatory drug treatment. Vaccine-related variables and exposure to different SARS-CoV-2 variants before or after vaccination, and consequently the diversity of pre-existing immunity of future vaccine recipients, differ notably between regions of the world and countries. This variability introduces uncertainty in extrapolating virus neutralisation data between countries. Therefore, to inform public health policies, it is necessary to characterise the anticipated sensitivity of emerging SARS-CoV-2 variants to vaccines in different populations using clearly defined cohorts.

The combination of Danish SARS-CoV-2 whole genome sequencing surveillance, vaccination registries and biobanking enables the selection of clearly defined cohorts. Of relevance, the cohorts selected in the present study included primarily older individuals, who represent the population groups most likely to receive the updated XBB.1.5 vaccines. An additional strength of the study includes the use of live, replication competent SARS-CoV-2 clinical isolates that represent the pathogens circulating in the population. A limitation of the study is the limited sample sizes; although, the relative neutralisation data are in agreement with the findings of others (2,5,6,8–10).

In conclusion, this study provides antibody cross-neutralisation data for the newly emerged Omicron subvariant BA.2.86 and globally dominating subvariant EG.5.1 in a population that received mRNA vaccines as primary regimen and boosters, and who experienced an Omicron BA.2 wave followed by smaller waves of BA.4/5 and a subsequent dominance of XBB.1 recombinant variants. We demonstrate that BA.2.86 neutralisation is comparable to that of XBB.1 and EG.5.1 in individuals who experienced XBB.1.5 breakthrough infection, BA.4/5 breakthrough infection, or BA.4/5 bivalent vaccination.

## Methods

### Ethics statement

This study is part of national infectious disease surveillance and response to an emerging virus variant performed on surplus biological material by Statens Serum Institut, a governmental institute under the Danish Ministry of Health, according to Section 222 of the Danish Health Act and following data protection regulations. The serum or plasma samples used were pseudonymised and described so that individuals cannot be identified. This study is therefore exempt from ethical review and did not require informed consent.

### Virus isolation

In a 24-well plate, 3.5 × 10^4^ of Vero E6/Vero/Vero-hSLAM/CaLu-3 cells or 7 × 10^4^ of iGROV-1 cells were seeded per well and cultured overnight, with the exception of CaLu-3 cells that were cultured for three days, in culture medium (Dulbecco’s Modified Eagle Medium [DMEM], 10% foetal calf serum and 1% Penicillin/Streptomycin) at 37°C, 5% CO_2_. All variants were isolated on Vero E6 except the B.1 isolate (isolated on Vero) and XBB.1.4.1 and BA.2.86 isolates (isolated on CaLu-3). Successful isolation of BA.2.86 was also obtained with iGROV-1 cells, but was not used for further characterisation here. Prior to inoculation, the monolayers were washed with 1 mL phosphate buffered saline (PBS). 150 µL infection media (DMEM and 1% Penicillin/Streptomycin) and 150 µL of SARS-CoV-2 PCR positive sample (throat/nasal swabs in PBS/Ringer’s solution, or tracheal secrete) were added to 80-90% confluent cell monolayers. The cells were incubated with the inoculum for 1 hour at 37°C, 5% CO_2_, followed by the addition of 1 mL growth media (DMEM, 1% Penicillin/Streptomycin, 1.5 µg/mL Amphotericin B, 10% foetal calf serum). The cell culture supernatant was harvested 72-96 hours post-inoculation, clarified by centrifugation at 300 × g for 5 min, and stored at -80°C. All cell culture reagents were from Gibco, ThermoFisher Scientific, Waltham, Massachusetts, USA.

### Virus propagation

To expand the virus stocks, 1.5 × 10^6^ Vero E6 cells were seeded in 75 cm^2^ flasks and incubated overnight at 37°C, 5% CO_2_. The cell monolayer was washed once with 5 mL PBS and inoculated with 50-100 µL Passage 1 virus diluted in 2 mL DMEM. After a 1-hour incubation with the inoculum at 37°C, 5% CO_2_, 10 mL virus growth media (DMEM with 5% foetal calf serum, 1% Penicillin/Streptomycin, and 10 mM HEPES buffer) was added and incubated at 37°C, 5% CO_2_. The virus was primarily harvested 96 hours later. The passage 2 virus stock was clarified by centrifugation at 300 × g for 5 minutes and the supernatant stored in single-use aliquots at -80°C. All experiments were performed with passage 2 stocks except for isolates D614G and BA.5, where passage 3 was used. Isolate B.1 was only propagated in Vero cells. All virus stocks used in functional antibody characterisation were sequenced to confirm identity, the absence of cell culture-derived mutations, and the presence of lineage-specific mutations in the spike protein.

### Human serum

No individuals were sampled for the purpose of this study. Pseudonymised surplus serum or plasma samples for this study were obtained from the Danish National Biobank. Persons with specific SARS-CoV-2 vaccine or infection histories were identified by converging the SARS-CoV-2 whole genome sequence databases, vaccine registries, and biobank database. The selection criteria were: 1) adults only; 2) either sex assigned at birth; 3) serum/plasma sample availability in the biobank 14 to 90 days after an XBB.1.5 infection, BA.4/5 infection, or Comirnaty Original/Omicron BA.4-5 bivalent mRNA vaccine as a fourth vaccination against COVID-19; 4) absence of a documented SARS-CoV-2 vaccination or infection in the 14 to 90 day period; 5) available vaccination history; and 6) avoid duplicate samples from the same person.

#### Samples from XBB.1.5 breakthrough infected

From 3,062 individuals with sequence-confirmed XBB.1.5 spike variants, 13 individuals met the abovementioned selection criteria. Twelve individuals had received three or more vaccinations prior to the XBB.1.5 infection and seven had earlier had a documented SARS-CoV-2 infection. The ratio of males-to-females was 6:7 and the median age was 59 years (range: 31-79). The serum samples had been collected 25 to 74 days after a positive SARS-CoV-2 test (median: 43 days).

#### Samples from BA.4/5 breakthrough infected

From 46,918 individuals with sequence-confirmed BA.4 or BA.5 infection in 2022, 329 individuals met the selection criteria from which 30 individuals were randomly selected for inclusion in the present study. Twenty-three individuals had received three or more vaccinations prior to the BA.4/5 infection, three individuals had received two vaccinations prior to infection and four were unvaccinated. None experienced a prior documented SARS-CoV-2 infection. The ratio of males-to-females was 9:21 and the median age was 59.5 years (range: 20-76). The serum samples had been collected 14 to 83 days after a positive SARS-CoV-2 test (median: 28 days).

#### Samples from bivalent BA.4/5 vaccinated

Six individuals who had received the Comirnaty Original/Omicron BA.4-5 bivalent mRNA vaccine as a fourth vaccination met the selection criteria. None had a prior documented SARS-CoV-2 infection. The ratio of males-to-females was 3:3 and the median age 53.5 years (range: 28-73). The serum samples had been collected 16 to 87 days after a positive SARS-CoV-2 test (median: 42 days).

### Microneutralisation test

A 2-fold serial dilution of heat-inactivated serum/plasma samples was mixed with 300 × TCID_50_ SARS-CoV-2 virus and the solution incubated for 1 hour at 37°C, 5% CO_2_. The diluted serum/plasma with virus was subsequently added to Vero E6 cells in a 96-well tissue culture plate, seeded with 10^4^ cells per well the day prior, and incubated at 37°C, 5% CO_2_. The following day, the inhibition of virus infection was measured in a standard ELISA targeting the SARS-CoV-2 nucleocapsid protein. Culture medium was removed from the infected Vero E6 cell monolayers and the cells were washed twice with 100 μL PBS. The cells were fixed with cold 80% (v/v) acetone in PBS for 10 minutes. Following three wash steps with wash buffer (PBS containing 1% (v/v) Triton-X100) for 30 seconds, 100 μL of a SARS-CoV-2 nucleocapsid protein mouse monoclonal antibody clone 7E1B (1:4000 dilution; Cat. # BSM-41414M, Bioss, Woburn, Massachusetts, USA) was added and incubated for 5 minutes on an orbital shaker (300 rpm) at room temperature and subsequently for 1 hour at 37°C. The plates were washed and incubated with 100 μL of a 1:10,000 diluted goat anti-mouse IgG (H+L) cross-adsorbed HRP conjugate antibody (Cat. # A16078; Invitrogen, Waltham, Massachusetts, USA) for 5 minutes on an orbital shaker (300 rpm) at room temperature and subsequently for 1 hour at 37°C. The plates were washed five times with wash buffer for 30 seconds, followed by three washes with deionized water. A 100 μL TMB One Substrate (Cat. # 4380, KemEnTec, Denmark) was added and incubated for 15 minutes. The reaction was stopped with H_2_SO_4_ and absorbance read at 450 nm using 620 nm as a reference on a FLUOstar Microplate Reader (BMG LABTECH, Germany). The validated microneutralisation assay has comparable performance to other live virus neutralisation assays established at different European laboratories.

### Statistics and computational analysis

Included on each microneutralisation plate were quadruplicate wells containing cells with 300 × TCID_50_ SARS-CoV-2 virus without serum (virus control) and quadruplicate wells containing cells with virus diluent only (cell control). The neutralisation antibody titer was determined for each serum sample as the interpolation of a four-parameter logistic regression curve with the 50% virus level cut-off calculated for each assay plate: [(mean OD of virus control wells) + (mean OD of cell control wells)]/2. The reciprocal serum dilution corresponding to that well is reported as the 50% serum neutralisation titre for that sample.

## Ethical statement

This study was conducted as part of the Danish COVID-19 surveillance. According to Danish law, ethical approval is not required.

## Acknowledgements

We thank all the colleagues involved in the work at SSI for their contributions, regional laboratories for their collaboration regarding sending samples and genomes, and DURABLE-network for support. We thank Dennis Christensen from Statens Serum Institut, Denmark, and Bjoern Meyer from Institut Pasteur, France, for providing cell lines.

## Supplementary Information

**Figure S1.**
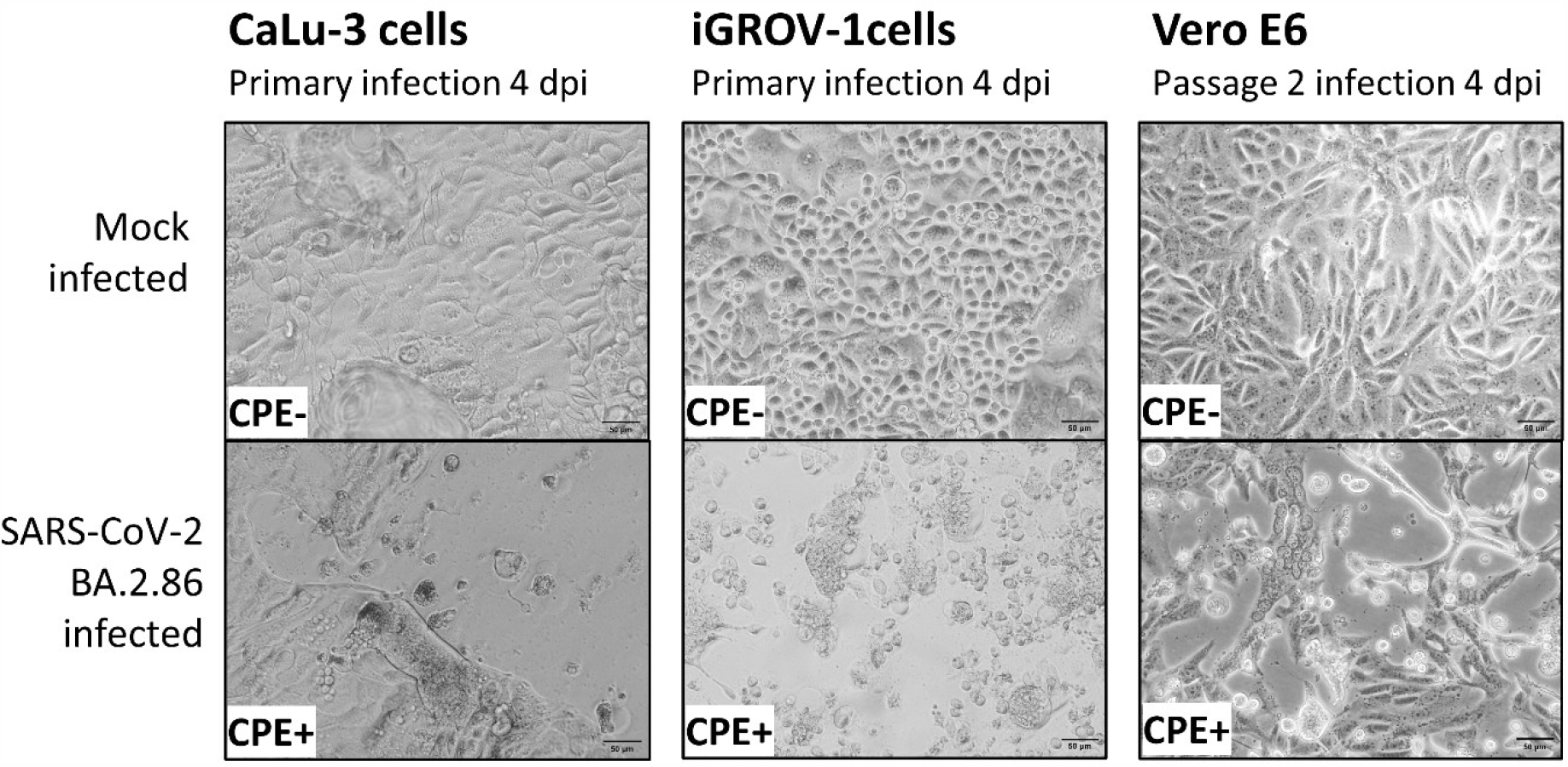
Cytopathic effect (CPE) of BA.2.86 induced on CaLu-3 cells and iGROV-1 cells four days post-inoculation with clinical sample material and Vero E6 cells four days post-inoculation with passage 1 virus produced in CaLu-3 cells. The passage 2 CPE induced in Vero E6 cells were similar when using CaLu-3 or iGROV-1 produced passage 1 virus.

## References

1. Rasmussen M, Møller FT, Gunalan V, Baig S, Bennedbæk M, Christiansen LE, et al. First cases of SARS-CoV-2 BA.2.86 in Denmark, 2023. Euro Surveill Bull Eur Sur Mal Transm Eur Commun Dis Bull. 2023;

2. Khan K, Lustig G, Reedoy K, Jule Z, Romer C, Karim F, et al. Evolution and neutralization escape of the SARS-CoV-2 BA.2.86 subvariant. medRxiv. 2023;

3. Uriu K, Ito J, Kosugi Y, Tanaka Y, Mugita Y, Guo Z, et al. Transmissibility, infectivity, and immune evasion of the SARS-CoV-2 BA.2.86 variant. Lancet Infect Dis Print. 2023;

4. Reeve L, Tessier E, Trindall A, Abdul Aziz NIB, Andrews N, Futschik M, et al. High attack rate in a large care home outbreak of SARS-CoV-2 BA.2.86, East of England, August 2023. Bd. 28, Eurosurveillance. 2023. s. 2300489.

5. Yang S, Yu Y, Jian F, Song W, Yisimayi A, Chen X, et al. Antigenicity and infectivity characterization of SARS-CoV-2 BA.2.86. Lancet Infect Dis Print. 2023;

6. Qu P, Xu K, Faraone JN, Goodarzi N, Zheng Y-M, Carlin C, et al. Immune Evasion, Infectivity, and Fusogenicity of SARS-CoV-2 Omicron BA.2.86 and FLip Variants. bioRxiv. 2023;

7. Aggarwal A, Fichter C, Milogiannakis V, Akerman A, Ison T, Silva MR, et al. TMPRSS2 activation of Omicron lineage Spike glycoproteins is regulated by TMPRSS2 cleavage of ACE2. bioRxiv. 2023;

8. Lasrado N, Collier AY, Hachmann N, Miller J, Rowe M, Schonberg E, et al. Neutralization Escape by SARS-CoV-2 Omicron Subvariant BA.2.86. bioRxiv. 2023;

9. Sheward D, Yang Y, Westerberg M, Öling S, Muschiol S, Sato K, et al. Sensitivity of BA.2.86 to prevailing neutralising antibody responses. Lancet Infect Dis Print. 2023;

10. An Y, Zhou X, Tao L, Xie H, Li D, Wang R, et al. SARS-CoV-2 Omicron BA.2.86: ess neutralization evasion compared to XBB sub-variants. bioRxiv. 2023

